# The Association between the JAK-STAT Pathway and Hypertension among Kenyan Women Diagnosed with Breast Cancer

**DOI:** 10.1101/2024.06.07.597892

**Authors:** John Gitau, Godfrey Kinyori, Shahin Sayed, Mohammad Saleem, Francis W Makokha, Annet Kirabo

## Abstract

**Background:** Breast cancer is the most common malignant tumor in women worldwide, and disproportionately affects Sub-Saharan Africa compared to high income countries. The global disease burden is growing, with Sub-Saharan Africa reporting majority of the cases. In Kenya, breast cancer is the most commonly diagnosed cancer, with an annual incidence of 7,243 new cases in 2022, representing 25.5% of all reported cancers in women. Evidence suggests that women receiving breast cancer treatment are at a greater risk of developing hypertension than women without breast cancer. Hypertension prevalence has been on the rise in SSA, with poor detection, treatment and control. The JAK-STAT signaling is activated in hormone receptor-positive breast tumors, leading to inflammation, cell proliferation, and treatment resistance in cancer cells. We sought to understand the association between the expression of JAK-STAT Pathway genes and hypertension among Kenyan women diagnosed with breast cancer.

**Methods:** Breast tumor and non-tumor tissues were acquired from patients with a pathologic diagnosis of invasive breast carcinoma. RNA was extracted from fresh frozen tumor and adjacent normal tissue samples of 23 participants who had at least 50% tumor after pathological examination, as well as their corresponding adjacent normal samples. Differentially expressed JAK-STAT genes between tumor and normal breast tissues were assessed using the DESEq2 R package. Pearson correlation was used to assess the correlation between differentially expressed JAK-STAT genes and participants’ blood pressure, heart rate, and body mass index (BMI).

**Results:** 11,868 genes were differentially expressed between breast tumor and non-tumor tissues. Eight JAK-STAT genes were significantly dysregulated (Log2FC ≥ 1.0 and an Padj ≤ 0.05), with two genes (CISH and SCNN1A) being upregulated. Six genes (TGFBR2, STAT5A, STAT5B, TGFRB3, SMAD9, and SOCS2) were downregulated. We identified STAT5A and SOCS2 genes to be significantly correlated with elevated systolic pressure and heart rate, respectively.

**Conclusions:** Our study provides insights underlying the molecular mechanisms of hypertension among Kenyan women diagnosed with breast cancer. Understanding these mechanisms may help develop targeted treatments that may improve health outcomes of Kenyan women diagnosed with breast cancer. Longitudinal studies with larger cohorts will be needed to validate our results.

## 1 Introduction

Breast cancer (BC) was the most common cancer in women in 157 countries out of 185 and accounted for 670,000 deaths globally in 2022 (Arnold *et al*., 2022a). Even though the disease incidence in Sub-Saharan Africa (SSA) is lower than in high-income countries, the region is disproportionately affected by breast cancer mortality (Martini *et al*., 2022),(Anyigba, Awandare and Paemka, 2021). In 2022, Low and middle-income countries (LMIC) registered more than half of the reported global breast cancer diagnoses and two-thirds of all breast cancer deaths (Arnold *et al*., 2022b; Wilkinson and Gathani, 2022). In Kenya, BC is a major public health concern, with an increasing incidence and mortality(Sung *et al*., 2021). According to the GLOBOCAN 2022 report, breast cancer was the second most prevalent cancer in Kenya and the most common cancer type among women, accounting for 25.5% of all new cancer types. Breast cancer incidence in Kenya has steadily increased over the last decade, with an estimated 2,296,840 new cases in 2022 and an estimated mortality of 66,103 women during the same year.

There is growing evidence of a bidirectional relationship between cardiovascular diseases and cancer (Bertero, Ameri and Maack, 2019; Cai *et al*., 2021; Guha *et al*., 2021; Chianca *et al*., 2022), whereby each can influence the development, progression, and treatment outcomes of the other. Mechanisms such as shared risk factors, systemic inflammation, oxidative stress, and endothelial dysfunction may influence this bidirectional relationship (Ganz *et al*., 2011; Wang *et al*., 2021). The regional heterogeneity in hypertension prevalence may be attributed to differences in the level of risk factors such as physical inactivity, alcohol consumption, high sodium intake, and low potassium intake (Mills, Stefanescu and He, 2020; Zhou *et al*., 2021; Vacca *et al*., 2023). Despite the rising prevalence of hypertension, particularly in SSA countries, there are few comprehensive assessments for hypertension early detection and treatment (Okello *et al*., 2020).

Breast cancer is associated with a chronic inflammatory state that can contribute to hypertension development (Jahan, Al-saigul and Abdelgadir, 2016; Ruan *et al*., 2023). Chronic inflammation is characterized by persistent immune system activation, which results in the production of pro-inflammatory cytokines such as interleukin-6 (IL-6) and tumor necrosis factor-alpha (TNF-IZ) (Jahan, Al-saigul and Abdelgadir, 2016; Ruan *et al*., 2023). Chemotherapy, radiation therapy, and hormonal therapy can all contribute to the development of a hypertensive phenotype (Darby *et al*., 2013; McGowan *et al*., 2017; Shin and Noh, 2018). Anthracyclines and taxanes, two chemotherapy drugs, have been linked to an increased risk of hypertension (de Jesus *et al*., 2022). Tamoxifen and aromatase inhibitors, for example, can cause hypertension by promoting vasoconstriction and endothelial dysfunction (Clarke *et al*., 2001; Blaes *et al*., 2019). Postmenopausal women with hypertension may have up to a 15% increased risk of developing breast cancer. This may be due to shared pathophysiological pathway mediated by adipose tissue, which may cause inflammation (Siiteri, 1987; Balkwill, Charles and Mantovani, 2005; Han *et al*., 2017)

The JAK/STAT signaling pathway is essential in many biological processes, including cell growth, differentiation, and survival. This pathway’s dysregulation has been linked to the development of breast cancer and hypertension (Roger *et al*., 2021; Ayele *et al*., 2022). In this study, we analyzed the expression profile of genes involved in hypertension to look for associations with blood pressure, heart rate, and body mass index (BMI) in Kenyan women diagnosed with and without breast cancer to determine their possible roles in either disease diagnosis, progression or treatment response, including hypertension.

Understanding these mechanisms may help develop targeted treatments that may improve health outcomes of Kenyan women diagnosed with breast cancer

## 3 Methods

### 3.1 Participant recruitment, sample collection, and Ethical consideration

The study Participants were selected from outpatient breast clinics at Aga Khan University Hospital-Nairobi (AKU) and AIC Kijabe Hospital (KAIC). The study recruited patients with a pathologic diagnosis of invasive breast carcinoma who were undergoing surgery and agreed to have their breast tumor and non-tumor tissue from the definitive surgical specimen genetically analyzed. Patients with benign breast disease were excluded from this study.

The AKU (2018/REC-80(v3)) and KAIC (KH IERC-02718/0036/2019) Institutional Scientific and Research Ethics Committees approved this study. The Ministry of Health in Kenya granted permission to transfer samples to the Laboratory of Human Carcinogenesis, National Cancer Institute (NCI), Bethesda, Maryland (MOH/ADM/1/1/81) for sequencing.

### 3.2 RNA Extraction and Sequencing

Fresh frozen tumors and adjacent normal tissue samples from 45 participants were shipped on dry ice to the National Cancer Institute’s Laboratory of Human Carcinogenesis in Bethesda, Maryland, for RNA extraction and sequencing. RNA extraction was performed on tumor samples from 23 participants who had at least 50% tumor composition, as well as their corresponding adjacent normal samples. Total RNA was isolated with the TRIzol reagent (Thermo Fisher Scientific) and treated with DNase.

The Agilent TapeStation High Sensitivity RNA ScreenTape was used to assess the integrity of the isolated RNA (Agilent Technologies). Total RNAseq was carried out at Leidos Biomedical Research, Inc.’s Sequencing Facility at the Frederick National Laboratory for Cancer Research. NEBNext UltraII Directional RNA Library Prep and paired-end sequencing were used on the NovaSeq S2 to sequence the samples.

### 3.3 RNAseq data pre-processing

The sequenced samples had an average of 57 million pass filter reads, with 93.7% of bases scoring higher than Q30. The samples’ reads were trimmed for adapters and low-quality bases with Cutadapt before being aligned with the reference genome (hg38) and annotated transcripts with STAR. All samples had a 96% mapping rate on average. Picard software was used to compute the mapping statistics. Picard’s Mark Duplicate utility was used to assess library complexity in terms of unique fragments in mapped reads. The samples contained up to 0.05% ribosomal bases and had 70% non-duplicate reads with 59% mRNA bases on average. All samples were aligned to the reference genome (hg38) and gene expression quantified using STAR/RSEM tools in NextFlow’s RNAseq pipeline (version 3.8) (Lataretu and Hölzer, 2020). The raw counts were normalized in R statistical software using the trimmed mean of M values (TMM) (Robinson and Oshlack, 2010).

### 3.4 Differential gene expression analysis

Differentially-expressed JAK-STAT genes (DEGs) between tumor and normal breast tissues were assessed using the DESEq2 R package (Robinson, McCarthy and Smyth, 2009). Prior to analyzing differential gene expression, Principal Component Analysis (PCA) was done to determine sample clustering using variance stabilizing transformation (vsd) normalization extracted with DESEq2 analysis in ggplot2 R package. The raw counts data were thereafter used as an input for DESEq2 analysis, with the cut-off criteria for assessing the DEGs set at |log2FC|>1.0 and adjusted P<0.05. The resultant volcano plot was depicted using the Enhanced volcano plot R package (Blighe, 2018). In the differential expression analysis, a multiple testing correction was performed using the Benjamini-Hochberg procedure to reduce the number of false positives, and a false discovery rate (FDR) <5% was considered to identify the significantly dysregulated pathways within each group sample group.

### 3.5 Correlational analysis

A Pearson’s correlation coefficient was used to examine differentially expressed JAK-STA genes using the packages corrplot, and psych in R. Systolic and diastolic blood pressure, heart rate, weight, and BMI were correlated with normalized counts of key JAK-STAT genes. A single data frame with clinical information parameters and normalized counts was formed, and thereafter their correlation was calculated.

## 4 Results

We collected and extracted RNA from twenty-three (23) tumor and normal paired samples, performed differential gene expression analysis, identified differentially expressed JAK-STAT genes, and performed a correlational analysis with patient clinical data. One sample was excluded from the study as it did not meet the quality threshold. The gene expression levels of the twenty-two (22) paired samples from tumor and non-tumor adjacent tissues were compared. A total of 11,868 differentially expressed genes from the initial 44,584 genes using an adjusted P-value of <0.05, and a Log Fc of +/-1 (Figure 3) were identified. Of the differentially expressed genes, eight (8) JAK-STAT genes were significantly expressed between normal and tumor tissues.

**Figure 1:**
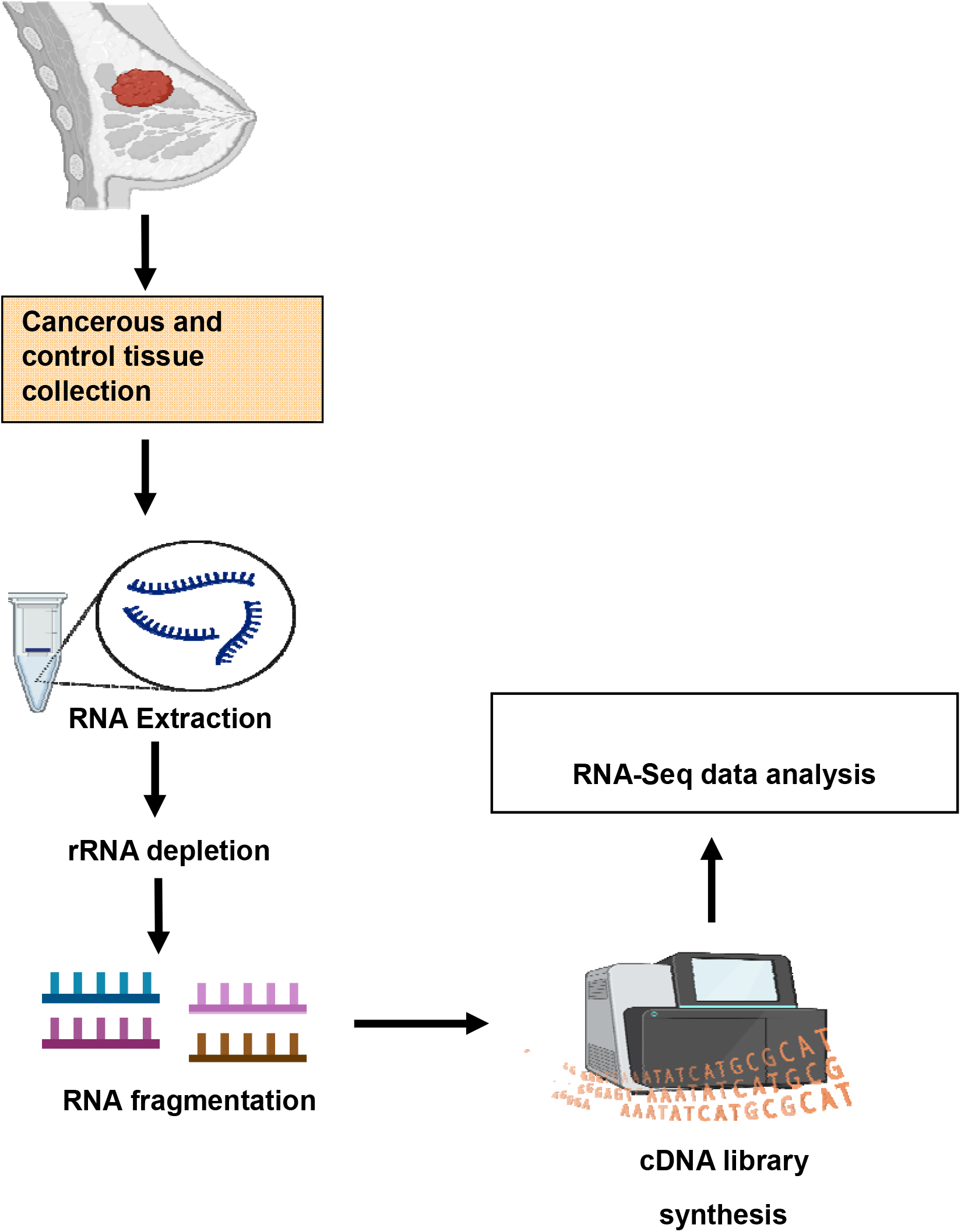
The schematic diagram shows the experimental design of isolation of human monocytes, followed by RNA extraction and performing bulk-RNA-seq.

Exploratory analysis using the Principal Component Analysis (PCA) in R showed sample clustering of the twenty-two paired breast tissues. Tumor and normal samples clustered separately as depicted in Figure 2. PCA plot visualized the overall variance and patterns present in the gene expression data, between tumor and normal samples. Additionally, the PCA plot checked for outliers in the dataset.

**Figure 2:**
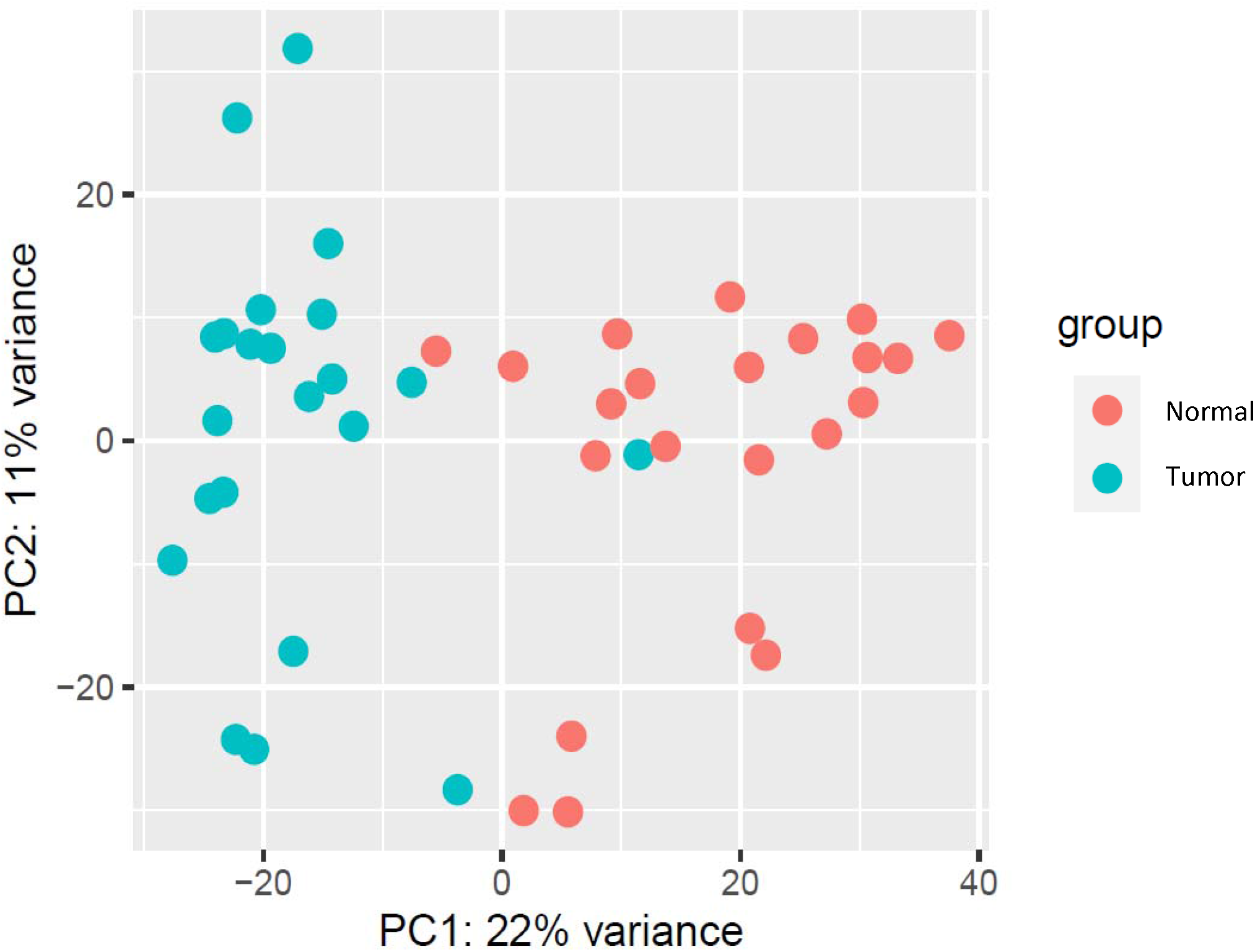
A PCA plot of normalized RNA-Seq count data. Each dot represents a sample, and twenty-two paired breast tumors and normal tissues were compared.

**Figure 3:**
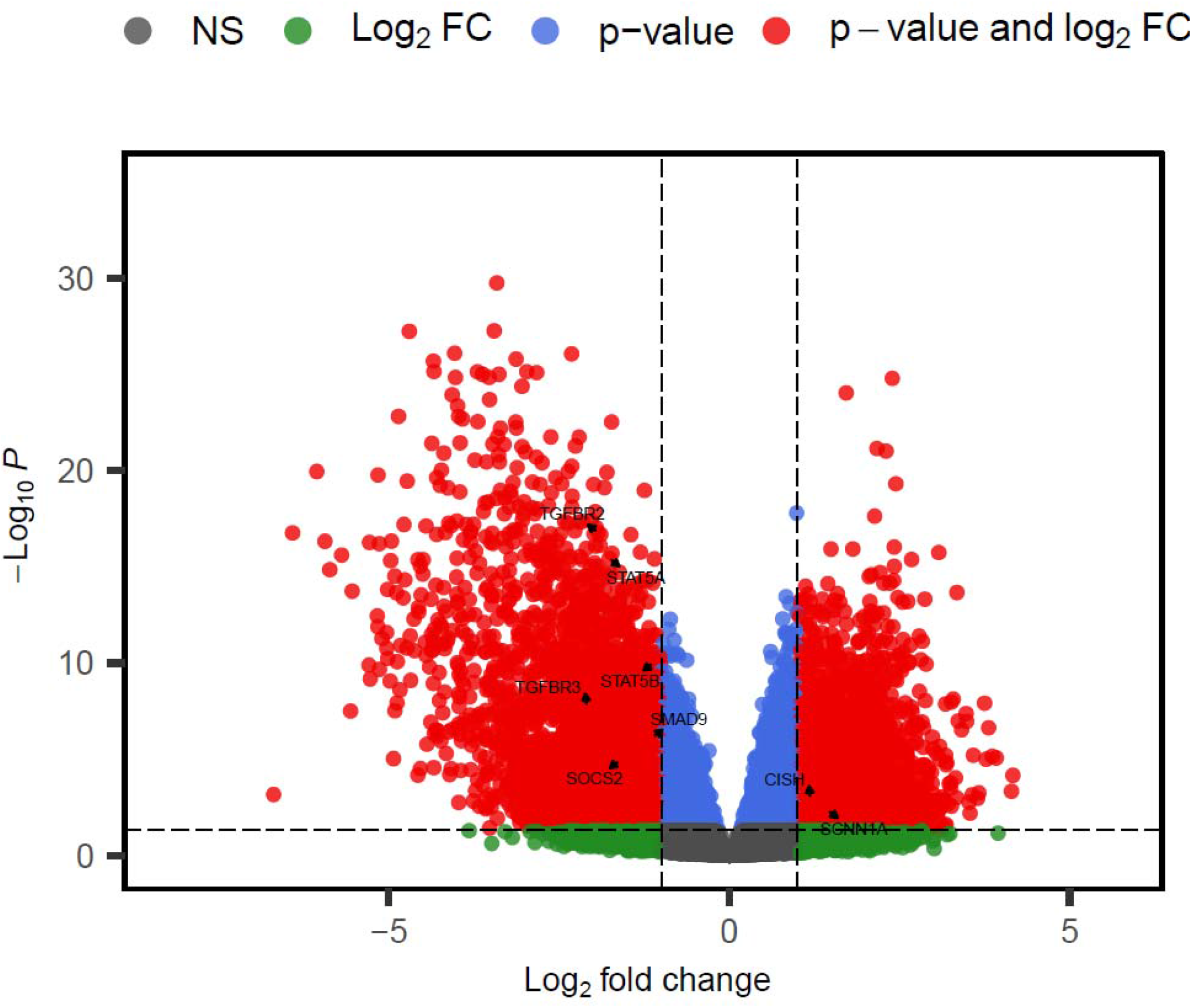
Volcano plot showing up and down differentially expressed JAK-STAT genes, between tumor and non-tumor adjacent tissues. Each dot represents a gene; blue dots represent genes with above cutoff adjusted P-values but with insignificant log2 fold change. Red dots represent significant genes, above an adjusted P-value and log2 fold change cutoff. Green dots represent genes with significant log2 fold change but below cutoff adjusted P-values. Grey dots represent non-significant genes (NS). A significance threshold of an absolute fold change of ≥ 1.0 and an adjusted P ≤0.05 were applied.

Eight out of the 64 previously characterized JAK-STAT genes were key in this study (Hu *et al*., 2021). Six (6) genes were upregulated while two (2) were downregulated. The downregulated genes included CISH and SCNN1A, while the downregulated genes included TGFBR2, STAT5A, STAT5B, TGFRB3, SMAD9 and SOC2 (table 1).

**Table 1:** Significantly expressed JAK-STAT genes with a log2 foldchange of more than +1 and less than -1, and an adjusted pvalue of <0.05. Seven genes were downregulated, while two genes were upregulated.

|  | Gene Name | log2FoldChange | Pvalue | Adjusted Pvalue (padj) |
| --- | --- | --- | --- | --- |
| 1 | TGFBR2 | -1.927096647 | 5.91E-20 | 1.64E-17 |
| 2 | STAT5A | -1.764815254 | 1.94E-18 | 3.63E-16 |
| 3 | STAT5B | -1.114759028 | 2.17E-12 | 9.58E-11 |
| 4 | TGFBR3 | -2.093546869 | 7.00E-10 | 1.73E-08 |
| 5 | SMAD9 | -1.130709869 | 4.81E-08 | 7.41E-07 |
| 6 | SOCS2 | -1.60570176 | 1.08E-06 | 1.14E-05 |
| 7 | CISH | 1.188695794 | 0.000197662 | 0.001082063 |
| 8 | SCNN1A | 1.437997521 | 0.000971912 | 0.004243666 |

**Table 2:** A table showing nineteen tumor samples with their corresponding Systolic blood pressure, diastolic blood pressure, heart rate, height, weight, and Body Mass Index (BMI).

| Sample id | Sample type | Systolic Pressure | Diastolic Pressure | heart rate | Height | Weight | BMI |
| --- | --- | --- | --- | --- | --- | --- | --- |
| Sample 1 | tumor | 120 | 78.5 | 83 | 163.5 | 101.5 | 38 |
| Sample 2 | tumor | 134 | 91 | 74 | 160 | 71 | 27.7 |
| Sample 3 | tumor | 144 | 94 | 88 | 157 | 63 | 25.6 |
| Sample 4 | tumor | 125 | 84.25 | 88.25 | 168 | 95 | 33.7 |
| Sample 5 | tumor | 124 | 55 | 78 | 159 | 63 | 24.9 |
| Sample 6 | tumor | 121.6 | 81 | 85.6 | 170 | 77 | 26.8 |
| Sample 7 | tumor | 133 | 88 | 71 | - | 90 | - |
| Sample 8 | tumor | 124.8 | 70.3 | 113.5 | 159 | 70.3 | 27.83 |
| Sample 9 | tumor | 132.6 | 80.6 | 92.6 | 163 | 69.1 | 25.5 |
| Sample 10 | tumor | 142 | 94 | 94 | 142 | 59 | 22.5 |
| Sample 11 | tumor | 101 | 64 | 75 | 148.99 | 70.66 | - |
| Sample 12 | tumor | 137 | 92 | 79 | - | - | - |
| Sample 13 | tumor | 172 | 85 | 71 | 147 | 50 | 23.14 |
| Sample 14 | tumor | 147 | 82 | 100 | 152 | 70 | 30.30 |
| Sample 15 | tumor | 150 | 87 | 64 | 164 | 84 | 31.42 |
| Sample 16 | tumor | 129 | 83 | 96 | 172 | 75.5 | 25.52 |
| Sample 17 | tumor | 116 | 60 | 86 | 160 | 74.4 | 29.06 |
| Sample 18 | tumor | 114 | 72 | 75 | - | 67.3 | - |
| Sample 19 | tumor | 113 | 67 | 98 | 162.5 | 73.8 | 27.95 |

The correlation analysis between clinical data from breast cancer patients and the expression levels of JAK-STAT genes revealed a positive correlation between the SOCS2 gene and heart rate (see Figure 4). Additionally, the expression of the STAT5A gene showed a positive correlation with systolic blood pressure.

**Figure 4:**
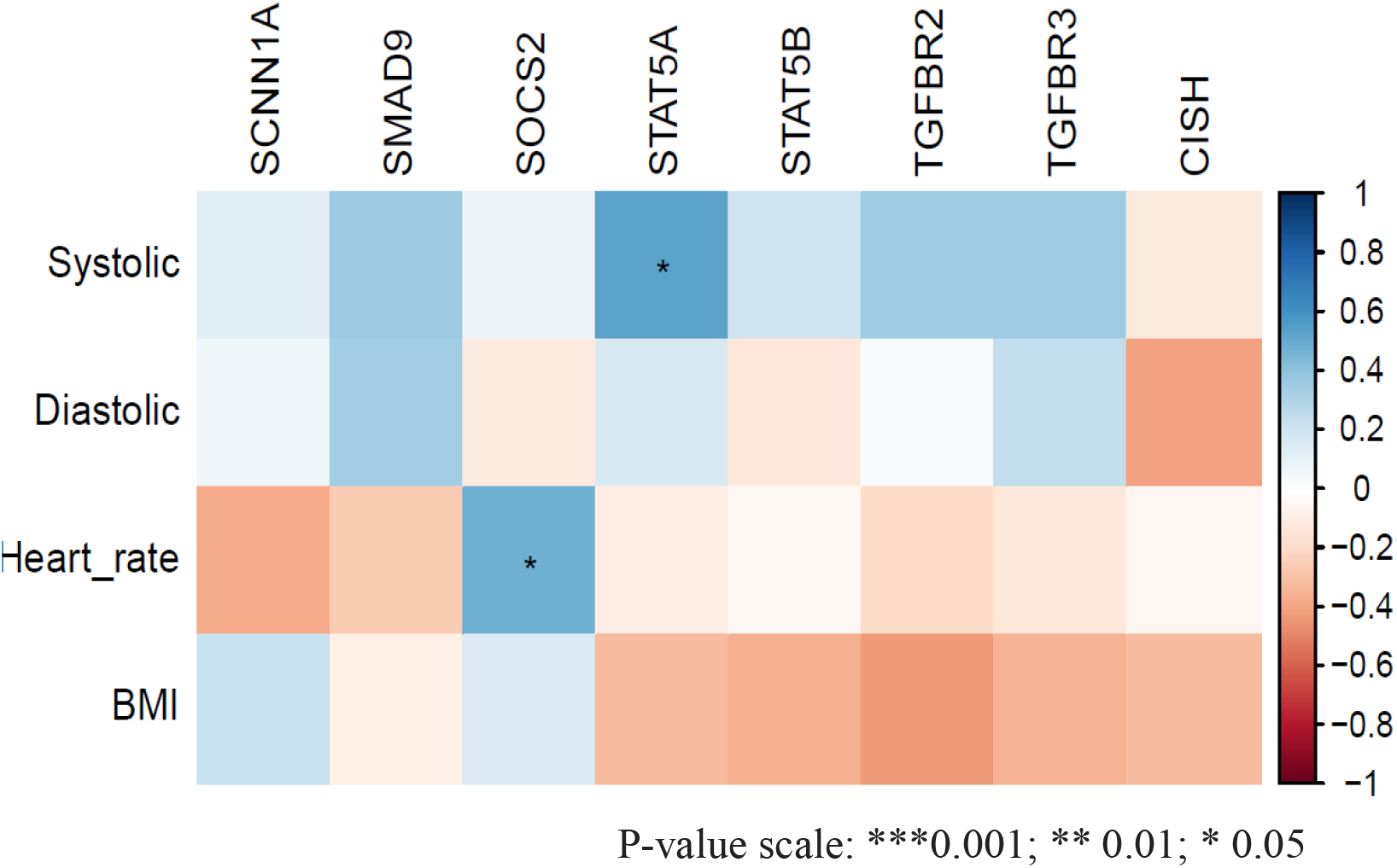
A correlational Matrix illustrating the correlation, and statistical significance between the JAK/STAT genes and key clinical characteristics of the Kenyan breast cancer patients. The SOCS2 and STAT5A genes are positively correlated with heart rate and systolic blood pressure with a P-value of 0.05. The color-coded correlations depict their direction and strength, as outlined in the legend on the right.

**Figure 5:**
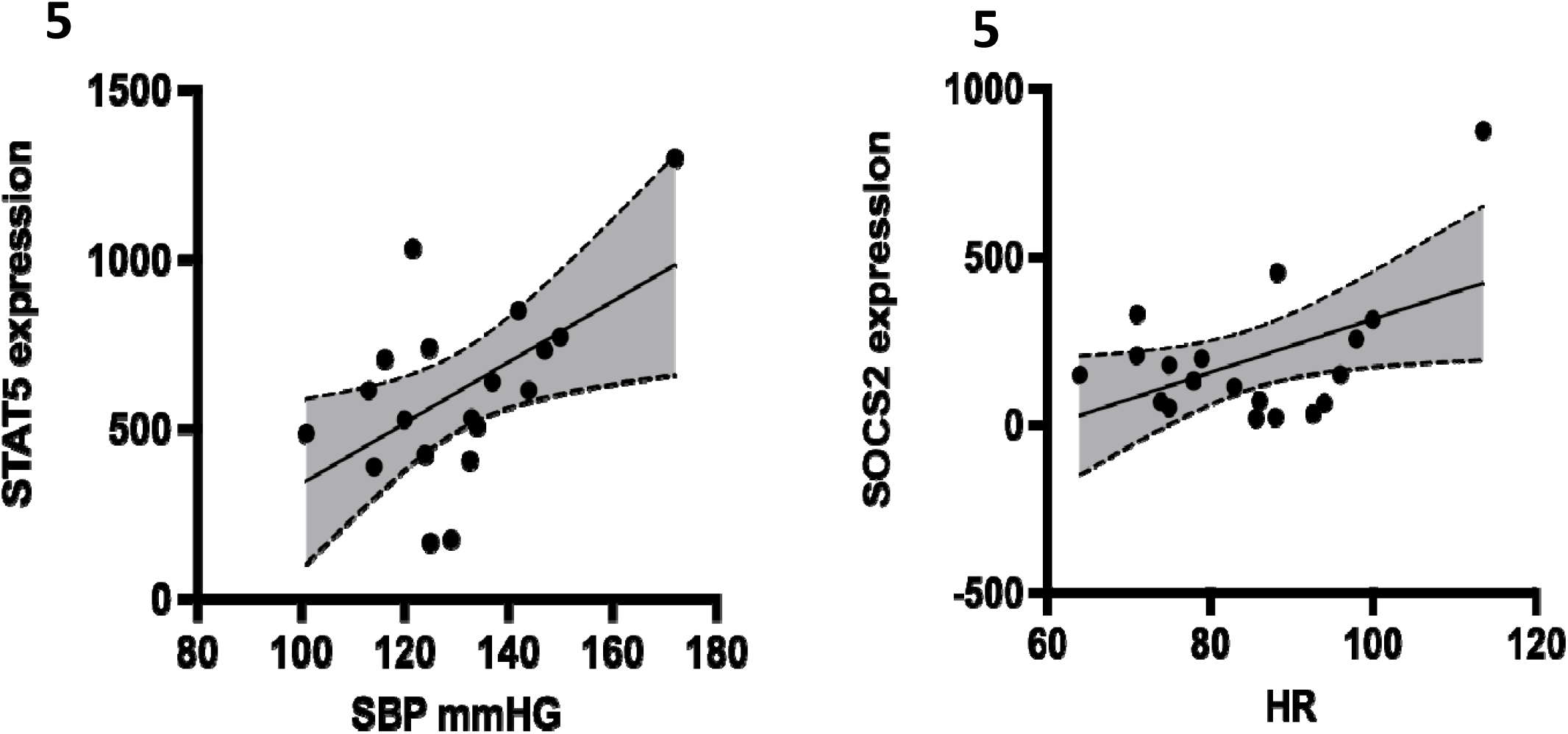
A diagrammatic representation of the positive correlation between the expression of STAT5A (**5A**), and SOCS2 (**5B**), genes and key clinical characteristics of Kenyan breast cancer patients. STAT5A gene is positively correlated with systolic blood pressure (SBP), while SOCS2 gene is positively correlated with heart rate (HR).

## 5 Discussion

In this study, we investigated the association between the expression of JAK-STAT genes with hypertension among Kenyan women diagnosed with breast cancer. We also investigated whether the expression of these genes is associated with clinical parameters such as blood pressure, heart rate, and BMI. Our findings indicate significant dysregulation in the expression of eight genes and our analysis showed that STAT5A and SOCS2 positively correlated with systolic blood pressure and heart rate respectively.

Our analysis identified a significant positive correlation between the expression of SOCS2 gene and heart rate. Elevated heart rate is associated with elevated blood pressure, increasing the risk of hypertension (Reule and Drawz, 2012). Several studies have reported a correlation between the downregulation of SOCS2 gene and breast cancer pathogenesis. A study by (Sutherland *et al*., 2004) observed that the expression of SOCS2 inhibits the growth of breast cancer cells. The investigators reported that aberrant methylation of SOCS2 gene correlates with transcriptional silencing of both breast and ovarian cell lines and that SOCS2 gene suppressed the growth of both tumor cells. We identified downregulation of the SOCS2 gene among Kenyan breast cancer patients. These findings were also in line with studies by (Farabegoli, 2005; Haffner *et al*., 2007). According to (Haffner *et al*., 2007), breast cancer patients with a higher expression of SOCS2 gene significantly lived longer than those with a lower expression profile. In our study, we observed the downregulation of SOCS2 gene, suggesting a more aggressive tumor among breast cancer Kenyan women. Between 2009 and 2019, diabetes associated deaths in Nairobi, the capital of Kenya increased by 65% (Manyara *et al*., 2024). Hypertension is twice as frequent in patients with diabetes compared with those who do not have diabetes (Petrie, Guzik and Touyz, 2018), due to salt and water retention. Studies have shown a decreased expression of SOCS2 gene in diabetic patients, and there exists a correlation between diabetes and hypertension; we suggest a that a downregulation of this gene in breast cancer patients may predispose them to a hypertensive phenotype.

According to the American heart Association, normal blood pressure readings are usually less than 120mmHg, with elevated blood pressure ranging between 120 and 129mmHg. High blood pressure (hypertension) is exhibited by systolic blood pressure reading of above 130mmHg (Ali *et al*., 2018). We observed that 19 breast cancer women were hypertensive. We also observed a significant correlation between the downregulation of STAT5A JAK-STAT gene and systolic blood pressure. STAT5A gene is predominantly in the mammary gland (Hennighausen and Robinson, 2008). There exist conflicting findings in literature pertaining to the role of STAT5A in breast cancer pathogenesis. On one hand, there are studies that associate the STAT5A with survival of breast cancer tissues due to the role of STAT5A in promoting breast tissue proliferation (Yamashita and Iwase, 2002a; Vafaizadeh *et al*., 2010). On the other hand, other studies associate expression of STAT5A gene with a better breast cancer prognosis, as the gene promotes mammary gland epithelial cell differentiation (Miyoshi *et al*., 2001; Yamashita and Iwase, 2002b; Wagner and Rui, 2008; Peck *et al*., 2012).

It is established that STAT5A gene is involved in various cellular processes, including cell proliferation and differentiation (Halim *et al*., 2020). STAT5A also regulates endothelial function and its downregulation may reduce the ability of blood vessels to dilate thereby creating internal pressure that may result to hypertension. Additionally, downregulation of STAT5A gene may lead to excessive smooth muscle proliferation, ultimately resulting in vascular remodeling and increased vascular resistance that contributes to hypertension.

While our study suggests a potential mechanism contributing to hypertension among Kenyan breast cancer patients, several limitations should be acknowledged. The relatively small sample size and cross-sectional nature of our study may limit the generalizability of our findings. Consequently, we propose longitudinal studies with larger cohorts, to validate our results and elucidate the underlying mechanisms further.

Our study may potentially enhance the understanding of molecular mechanisms underlying hypertension among Kenyan women diagnosed with breast cancer. By studying the dysregulated JAK-STAT genes, findings demonstrate a possible synergistic and/or antagonistic molecular mechanisms associated with the dysregulation of JAK-STAT genes. Our study therefore highlights the need for further research using larger samples, to potentially decipher mechanisms involved between systolic blood pressure, breast cancer pathogenesis and hypertension.

## Acknowledgement

We thank the breast cancer patients at The Aga Khan hospital Nairobi, and AIC Kijabe mission hospital for allowing us to use their tissue samples for this work.

## Data accessibility

The RNA-seq data analyzed in this study is publicly accessible at the GEO database under Accession number: GSE225846. All other datasets are included in the article.

## Author’s contribution

FM conceived the initial idea, designed the research project, collected tumor and normal samples and set up experiments. JG wrote the manuscript drafts and performed data analysis. GK performed data analysis and reviewed the manuscript. SS, MS, FM, and AK read and reviewed the manuscript drafts, and provided supervision. All authors contributed to the revision and final editing of the manuscript.

